# *C. elegans* nucleolar RG repeats are sufficient for nucleolar accumulation but insufficient for sub-nucleolar compartmentalization

**DOI:** 10.1101/2024.12.19.629445

**Authors:** Emily L. Spaulding, Dustin L. Updike

## Abstract

Intrinsically disordered arginine-glycine (RG) repeat domains are enriched in multilayered biomolecular condensates such as the nucleolus. *C. elegans* nucleolar RG repeats are dispensable for nucleolar accumulation and instead contribute to the organization of sub-nucleolar compartments. The sufficiency of RG repeats to facilitate sub-nucleolar compartmentalization is unclear. In this study, we drive expression of full-length RG repeats in the *C. elegans* germline to test their ability to localize to nucleoli and organize into nucleolar sub-compartments *in vivo*. We find that repeats accumulate within germ cell nucleoli but do not enrich in the correct sub-compartment. Our results suggest RG repeats may indirectly influence nucleolar organization by creating an environment favorable for sub-nucleolar compartmentalization of proteins primarily based on their function within the nucleolus.

## INTRODUCTION

The organization of functionally related proteins and nucleic acids into biomolecular condensates (BMCs) has emerged as a central theme in cell biology. Dozens of unique BMCs have been described, ranging in composition from one to hundreds of proteins (Rostam et al., 2023). Some, such as nucleoli and germ granules, are comprised of multiple, immiscible compartments (Brangwynne et al., 2009, 2011; Phillips et al., 2012; Feric et al., 2016; Banani et al., 2017; Wan et al., 2018; Marnik and Updike, 2019; Manage et al., 2020). The mechanisms by which proteins are targeted to the correct BMC and organized into sub-compartments are unclear, especially in the cells of living animals.

Many BMCs form through condensation of proteins with intrinsically disordered regions (IDRs). IDRs lack a stable 3-dimensional structure and shuffle through thousands of conformations, making it challenging to characterize their contribution to BMC dynamics (Holehouse and Kragelund, 2023). Although IDRs are enriched in BMC proteins, IDRs are not necessarily determinants of condensation and their presence is not always required for targeting of a protein to the correct BMC (Girard et al., 1994; Snaar et al., 2000; Marnik et al., 2019; Martin and Holehouse, 2020; Spaulding et al., 2022).

One type of IDR consists of arginine-glycine (RG) repeats. RG repeats are enriched in many BMCs, including nucleoli, germ granules, and stress granules (Thandapani et al., 2013). RG repeats bind to RNA and form both homotypic and heterotypic interactions with other IDRs through electrostatic interactions, hydrogen bonding, and π-stacking (Hanakahi et al., 1999; Takahama et al., 2011; Chong et al., 2018). The strength of these interactions increases with multivalency. Many RG sequences include regularly spaced aromatic residues, such as tyrosine and phenylalanine, which contribute directly to condensation (Lin et al., 2017; Wang et al., 2018).

The complex *in vivo* environment influences condensation and BMC dynamics (Kim et al., 2023). In this study, we take advantage of the visually accessible and well-characterized *C. elegans* germline to study RG repeat function in a living animal (Figure 1A). *C. elegans* germ cells reside in a shared cytoplasm, with each nucleus surrounded by a nuclear membrane spotted with germ granules. Germ granules sit on top of nuclear pores, where they are sites of small RNA production and post-transcriptional gene regulation (Phillips and Updike, 2022) (Figure 1B). Inside each germ cell nucleus is a 2-3 micron-wide nucleolus. Nucleoli of mammals and other complex eukaryotes are partitioned into 3-5 sub-compartments that are thought to correspond to specific stages of ribosome biogenesis (Feric et al., 2016; Stenström et al., 2020; Shan et al., 2023). Nucleoli of lower eukaryotes, such as yeast and *C. elegans*, appear to contain only 2 compartments: (1) the fibrillar center (FC) where rRNA is transcribed and chemically modified and (2) the granular component (GC) where ribosomal subunits are assembled (Lafontaine et al., 2020; Spaulding et al., 2022; Tartakoff et al., 2022). The RG repeat-containing proteins Fibrillarin (FIB-1) and GAR1 (GARR-1) are found in the FC, where they function in rRNA methylation and pseudouridylation, respectively (Tollervey et al., 1993; Bousquet-Antonelli et al., 1997). The RG repeat-containing protein Nucleolin (NUCL-1) is found in the GC where it functions in several aspects of ribosomal processing (Bouvet et al., 1998; Ginisty et al., 1998; Roger et al., 2003; Rickards et al., 2007). Approximately 50% of nucleoli in the adult germline contain nucleolar vacuoles, which are void of nucleolar components (Xu et al., 2023) (Figure 1B,C).

**Figure 1:**
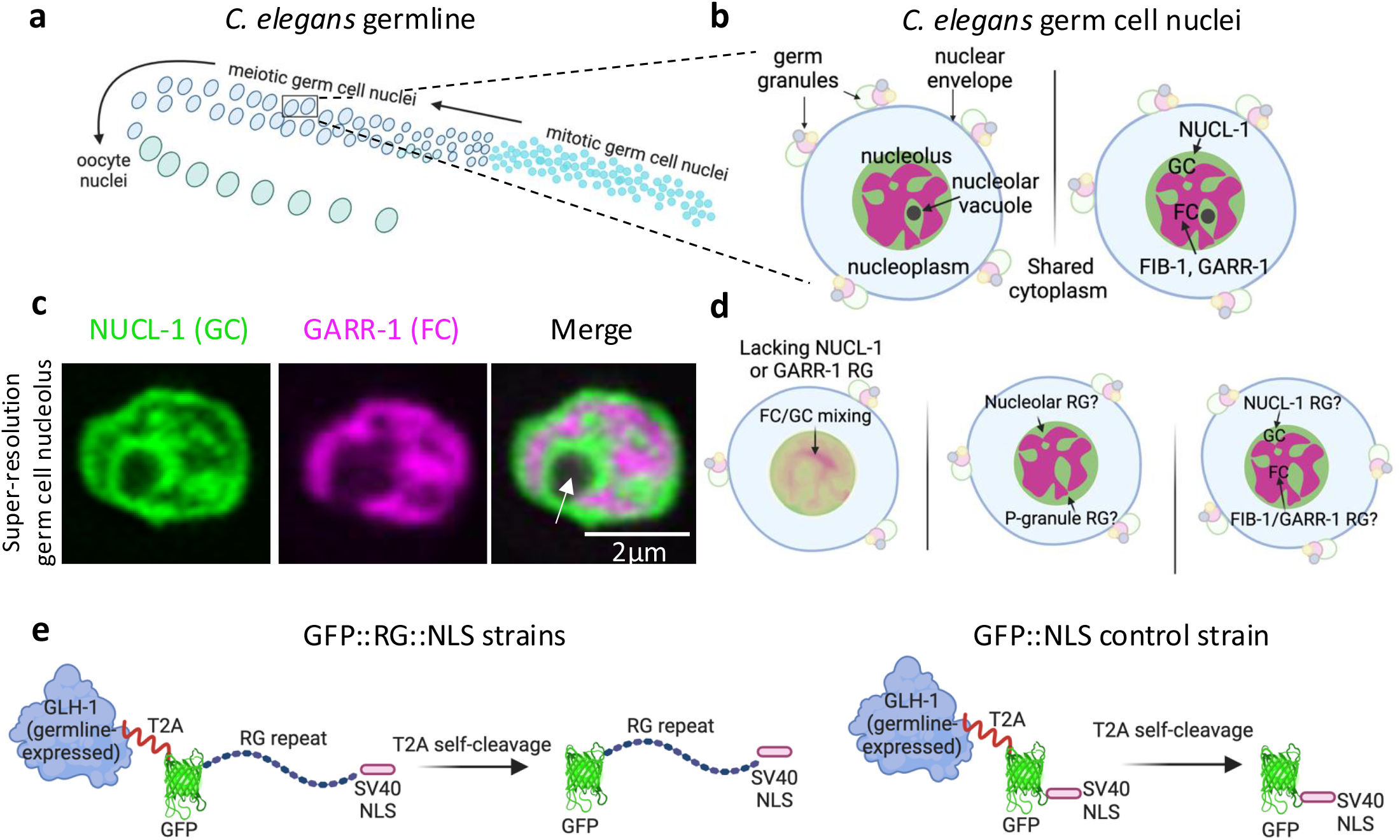
Studying RG repeat function in the *C. elegans* germline. **a** The adult hermaphrodite germline contains a progression of mitotic to meiotic germ cells that mature into oocytes ready for fertilization. **b** Schematic of 2 *C. elegans* pachytene germ cells. **c** Live, super-resolution (AiryScan) image of a germ cell nucleolus labeled with NUCL-1::GFP and GARR-1::wrmScarlet. White arrow points to a vacuole. Image is 1 plane of a 23-slice Z stack, cropped to focus on 1 nucleolus. **d** Deleting RG repeats causes FC/GC mixing (left-most nucleolus). Are RG repeats sufficient for nucleolar accumulation (center nucleolus)? Are RG repeats sufficient for sub-nucleolar compartmentalization (right-most nucleolus)? **e** Driver system for germline expression of RG repeats and GFP control.

We recently cataloged all RG repeats in *C. elegans* and found that nucleolar and germ granule repeats contain distinctive phenylalanine and tyrosine-rich consensus motifs, respectively. Despite the presence of a nucleolar phenylalanine-rich motif, RG repeats are dispensable for localization of proteins to the nucleolus. Instead, repeats organize proteins into nucleolar sub-compartments, and their removal leads to FC/GC mixing (Figure 1D) (Spaulding et al., 2022). How RG repeats direct sub-nucleolar compartmentalization is unclear, although recent work demonstrates they are not sufficient for compartmentalization in *Xenopus* oocytes (Lavering et al., 2023). In this study we drive expression of full-length nucleolar and germ granule RG repeats in the *C. elegans* germline and perform super-resolution imaging of living worms to test (1) if repeats independently accumulate in nucleoli and (2) if nucleolar repeats contain an intrinsic ability to organize into the correct sub-nucleolar domain (Figure 1D).

## RESULTS

### Driving RG repeat expression in the *C*. *elegans* germline

To visualize RG repeats exclusively within the *C. elegans* germline, we modified a system in which endogenous GLH-1 (Vasa) drives expression of a super-folder GFP (sGFP) (Goudeau et al., 2021). GLH-1 is one of the most highly expressed genes in the germline and its use as a driver promotes abundant and uniform expression of RG repeats (Campbell and Updike, 2015). In this system GLH-1 is separated from sGFP by a self-cleaving T2A peptide (GLH-1::T2A::sGFP) and GLH-1 expression is left intact. CRISPR/Cas9 genome editing was used to place full-length RG repeats followed by the SV40 nuclear localization signal (NLS) directly after sGFP in this strain. Upon T2A self-cleavage in the resulting strain (GLH-1::T2A::sGFP::RG::NLS), the sGFP-tagged RG repeat with NLS is released from GLH-1, resulting in fluorescently-tagged RG repeats in the nuclei of germ cells. As a control for possible GFP nucleolar enrichment and compartmentalization, sGFP with only the SV40 NLS was also included in each experiment (GLH-1::T2A::sGFP::NLS) (abbreviated GFP::NLS) (Figure 1E). Due to incomplete T2A cleavage from germ granule localized GLH-1, we expect to observe some residual GFP signal in germ granules within all strains using the GLH-1 driver system.

We inserted full-length RG repeat domains from three nucleolar proteins, NUCL-1, FIB-1, and GARR-1, into the expression system. The NUCL-1 repeat is the longest nucleolar RG repeat in *C. elegans* at 176 amino acids in length. The FIB-1 repeat is the third longest nucleolar repeat at 106 amino acids in length. Both repeats contain a phenylalanine-rich nucleolar consensus motif (“FRGGDRGGFR”) (Spaulding et al., 2022). The GARR-1 protein has two RG repeat domains, one at the N- and one at the C-terminus. The N-terminal 48 amino acid-long domain is phenylalanine-rich but does not contain the nucleolar consensus motif. The C-terminal 70 amino acid-long domain does contain the motif. We included the N-terminal RG repeat in this study to test if a nucleolar repeat without the consensus motif can accumulate and compartmentalize within nucleoli (Supplemental Figure 1A).

RG repeats from *C. elegans* germ granule proteins contain a tyrosine-rich consensus motif (“RGGRGGYRGGD”) (Spaulding et al., 2022). To test if an RG repeat with a germ granule motif will accumulate in nucleoli, we inserted the 53 amino acid-long PGL-1 RG repeat into the expression system (Supplemental Figure 1A). To mark nucleoli, wrmScarlet was placed on the C-terminus of endogenous GARR-1 (GARR-1::wSc) in each GFP::RG::NLS strain and the GFP::NLS strain. Finally, in a strain containing wrmScarlet-tagged GARR-1, we also placed sGFP at the C terminus of endogenous NUCL-1 (NUCL-1::GFP) to provide examples of nucleoli with correctly partitioned FC and GC sub-compartments. Western blots confirmed the expected size of each RG repeat insert. GFP::GARR-1 RG::NLS and GFP::PGL-1 RG::NLS repeats are expressed at the same level as GFP::NLS, but GFP::NUCL-1 RG::NLS and GFP::FIB-1 RG::NLS are expressed at lower levels (Supplemental Figure 1B,C). Evolutionarily conserved strict regulation of the synthesis and degradation of proteins with large IDRs has been demonstrated in organisms from yeast to humans (Gsponer et al., 2008). As the same endogenous driver is used for expression, lower levels of NUCL-1 and FIB-1 RGs suggest there may be cellular mechanisms in place to limit over-expression of the longest IDRs.

### RG repeats accumulate in germ cell nucleoli

Because RG repeats are dispensable for nucleolar accumulation of proteins in *C. elegans* germ cells, we asked if they are sufficient for accumulation. To visualize localization of RG repeats, we performed confocal imaging of live, adult worms. Endogenously-labeled GARR-1 and NUCL-1 proteins accumulate in germ cell nucleoli throughout the entire germline (Supplemental Figure 2A). The expected background GFP signal is visible in germ granules in all strains using the GLH-1 driver system, including the control GFP::NLS strain. This signal is a useful marker that defines the nuclear boundary (Supplemental Figure 2B, white arrow). GFP::NLS is observed throughout the nuclei of germ cells and a slight accumulation is visible in some nucleoli (Supplemental Figure 2B). In contrast, GFP-labeled RG repeats from NUCL-1, FIB-1, GARR-1, and PGL-1 are enriched in all germ cell nucleoli (Supplemental Figure 2C-F).

To measure nucleolar enrichment we performed super-resolution confocal imaging of meiotic, pachytene-stage germ cells and calculated the ratio of mean nucleolar fluorescence intensity to mean nucleoplasmic fluorescence intensity for at least 50 cells per strain. Endogenous GARR-1 and NUCL-1 demonstrated high nucleolar enrichment, as expected (Figure 2A). In contrast, GFP::NLS demonstrates a nucleolar enrichment of close to zero, which is significantly lower than endogenous GARR-1 in the same germ cells (Figure 2B). This result confirms that the GFP tag alone does not contribute to nucleolar enrichment in the GFP::RG repeat strains. The NUCL-1, FIB-1, GARR-1, and PGL-1 RG repeats all enrich in nucleoli, but with decreased efficiency compared to endogenous GARR-1 in the same cells (Figure 2C-F).

**Figure 2:**
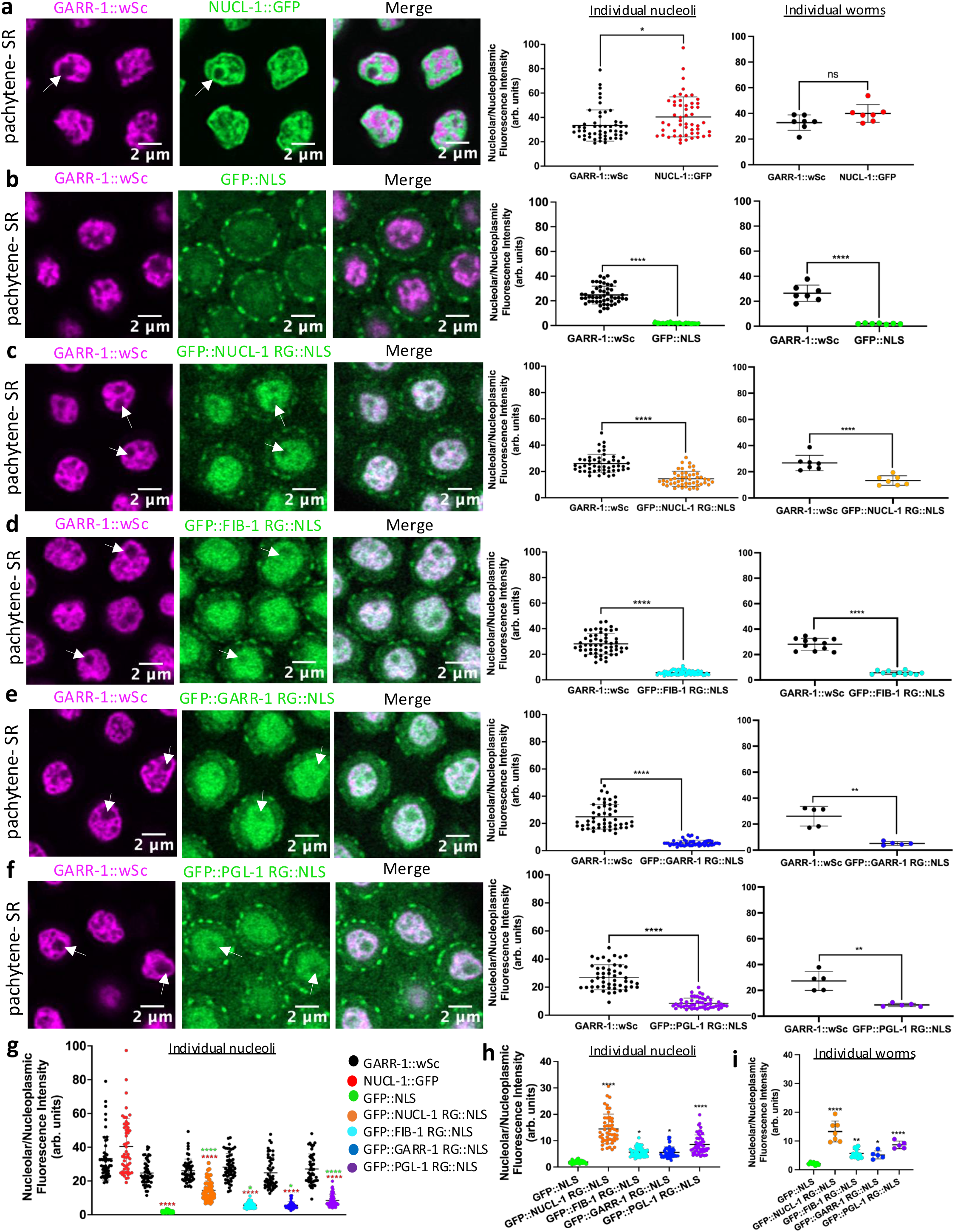
Nucleolar enrichment of RG repeats. **a-f** Single plane, super-resolution confocal images of 3-4 pachytene nucleoli and nucleolar enrichment quantification. **a** n=53 nucleoli from 7 worms (same image used in Fig 1c). **b-c** n=50 nucleoli from 7 worms. **d** n=55 nucleoli from 11 worms. **e-f** n=50 nucleoli from 5 worms. **a-f** Points on the left graph are individual nucleoli with means ± SD, points on the right graph are individual worms with means ± SD. **g** Comparison of nucleolar enrichment across all strains. Red asterisks compare against NUCL-1::GFP and green asterisks against GFP::NLS. **h-i** Comparison of RG repeat nucleolar enrichment against GFP::NLS, showing individual nucleoli (**h**) and individual worms (**i**). Data in **g-i** are the same data presented in **a-f. a-f** Unpaired t test with Welch’s correction. **g-i** One-way ANOVA with multiple comparisons. *p<.05, **p<.01, ****p<.0001.

Directly comparing the nucleolar:nucleoplasmic fluorescence intensity ratios of all strains confirms that each RG repeat accumulates in nucleoli less efficiently than GFP-tagged NUCL-1 or wrmScarlet-tagged GARR-1 (Figure 2G), but more efficiently than GFP::NLS (Figure 2H).

The NUCL-1 RG repeat is the longest repeat and displays the greatest nucleolar enrichment, most likely due to its high multivalence and capacity to engage in condensation-promoting molecular interactions. The germ granule PGL-1 RG repeat shows the second-highest nucleolar enrichment, despite being shorter than the FIB-1 RG repeat. This result may point to the stronger contribution of tyrosine residues to condensation compared to phenylalanine (Bremer *et al*., 2022). The GARR-1 repeat is phenylalanine-rich but does not contain the nucleolar consensus motif. Regardless, the GARR-1 repeat enriches in nucleoli, suggesting that non-motif phenylalanines in the GARR-1 repeat contribute to condensation behavior in a manner similarly to the consensus motif. In summary, nucleolar repeats both with (NUCL-1 and FIB-1) and without (GARR-1) the consensus motif and ranging in size from 176 to 48 amino acids accumulate in germ cell nucleoli. In addition, the germ granule PGL-1 RG repeat with a tyrosine-rich consensus motif accumulates in germ cell nucleoli.

### RG repeats do not display large-scale sub-nucleolar compartmentalization

Because RG repeats are required for enrichment of nucleolar proteins into the FC and GC, we asked if they are sufficient for this enrichment. To measure sub-nucleolar compartmentalization of RG repeats, we again used the super-resolution images of pachytene germ cells. The coefficient of variation (CV) is a measure of fluorescence variation and was calculated by dividing the standard deviation by the mean intensity of GFP fluorescence in individual nucleoli. In each nucleolus, the CV was also measured for endogenous GARR-1 as a positive control for compartmentalization. A high CV indicates heterogeneous distribution and more compartmentalization, while a low CV indicates homogeneous distribution and less compartmentalization (Spaulding et al., 2022).

In the strain with endogenously-tagged NUCL-1 and GARR-1, NUCL-1 enriches in the GC and GARR-1 enriches in the FC, as expected. NUCL-1 displays a slightly higher CV compared to GARR-1, most likely because the larger GC contains more areas of enrichment and depletion throughout nucleoli (Figure 3A). As expected, GFP::NLS shows no compartmentalization within nucleoli and has a significantly lower CV than endogenous GARR-1 (Figure 3B). NUCL-1, FIB-1, and GARR-1 RG repeats do not display large-scale compartmentalization and have significantly lower CVs than endogenous GARR-1 (Figure 3C-E). Endogenous NUCL-1 protein is restricted to the GC, but the NUCL-1 RG repeat localizes throughout the entire nucleolus and its expression overlaps with endogenous GARR-1 in the FC (Figure 3C, white arrows). Endogenous FIB-1 and GARR-1 proteins are restricted to the FC, but both FIB-1 and GARR-1 RG repeat expression spills over into areas of endogenous GARR-1 depletion that correspond to the GC (Figure 3D,E white arrows). In some nucleoli, RG repeat expression is lightest in spots that also lack endogenous GARR-1 expression, areas which most likely correspond to nucleolar vacuoles (Figure 3D-F, pink arrows) (Spaulding et al., 2022; Xu et al., 2023). As a native component of germ granules, the PGL-1 RG repeat was not expected to enrich in a specific nucleolar sub-compartment and shows significantly decreased CV compared to endogenous GARR-1 (Figure 3F). Although RG repeats have significantly lower CVs compared to endogenous GARR-1 or NUCL-1 (Figure 3G), they do have significantly higher CVs than GFP::NLS, suggesting some micro-organization within nucleoli (Figure 3H).

**Figure 3:**
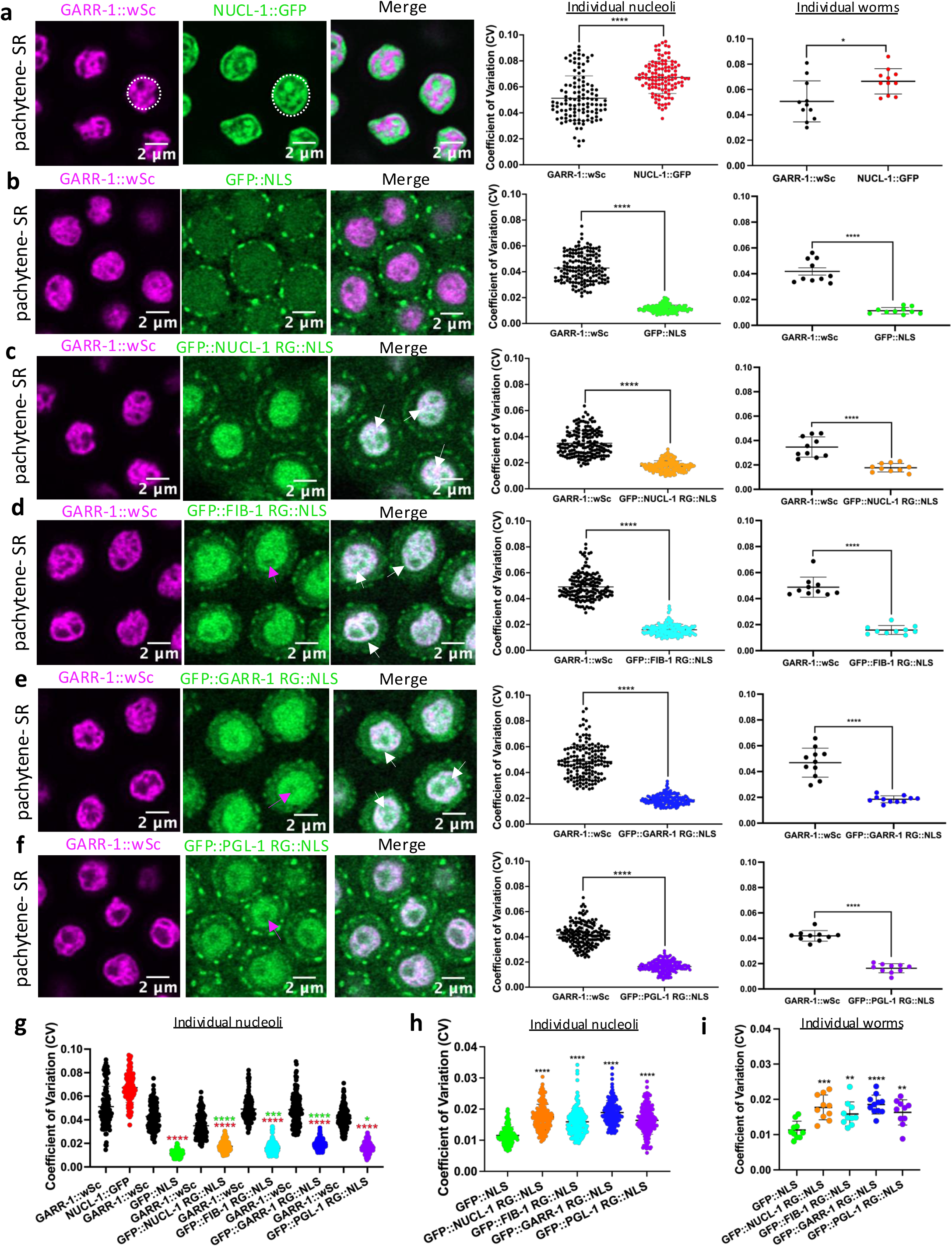
RG repeats do not display large-scale sub-nucleolar compartmentalization. **a-f** Single plane, super-resolution confocal images of 4-5 pachytene nucleoli and CV quantification. The white-dashed circle in (**a)** is an example of the area measured for CV. **a** n= 114 nucleoli from 11 worms. **b** n=153 nucleoli from 10 worms. **c** n=170 nucleoli from 10 worms. **d** n=159 nucleoli from 10 worms. **e** n=153 nucleoli from 11 worms. **f** n=175 nucleoli from 10 worms. **a-f** Points on the left graph are individual nucleoli with means**± SD, points on the right graph are individual worms with means ± SD. **g** Comparison of nucleolar enrichment across all strains. Red asterisks are against NUCL-1::GFP and green asterisks are against GFP::NLS. **h-i** Comparison of RG repeat strains against GFP::NLS, showing individual nucleoli (**h**) and individual worms (**i**). Data in **g-i** are the same data presented in **a-f. a-f** Unpaired t test with Welch’s correction. **g-i** One-way ANOVA with multiple comparisons. *p<.05, **p<.01, ***p<.001, ****p<.0001.

As a second way to measure RG repeat organization, we measured the Mander’s Colocalization Coefficient (MCC) for wSc and GFP using the same images used for CV analysis. MCC measures the fraction of one fluorophore that colocalizes with another. The Mander’s Overlap Coefficient 1 (M1) reports on the fraction of wSc fluorescence in areas with GFP fluorescence. The larger M1, the stronger the evidence for colocalization. Confirming the CV analysis, the overlap of GARR-1::wSc with GFP::NLS, NUCL-1RG, FIB-1RG, GARR-1RG, and PGL-1RG is significanctly higher than with endogenous NUCL-1 (Supplemental Figure 3).

As a third way to measure RG repeat compartmentalization, we created profile plots of mean wrmScarlet and GFP fluorescence across at least 50 pachytene nucleoli from each strain. The large standard deviation of endogenous NUCL-1 and GARR-1 fluorescence demonstrates the heterogeneous distribution of NUCL-1 in the GC and GARR-1 in the FC (Supplemental Figure 4A). In contrast, GFP::NLS and GFP-tagged RG repeats display smaller standard deviations across nucleoli, indicating more homogeneous distribution and a lack of precise compartmentalization (Supplemental Figure 4B-F). In summary, although nucleolar RG repeats are required for sub-nucleolar organization, they are insufficient to recognize and enrich in the appropriate nucleolar sub-compartment.

## DISCUSSION

RG repeat-containing nucleolar proteins such as Nucleolin, Fibrillarin, and GAR1 are precisely partitioned into immiscible sub-nucleolar compartments (Figure 4A). In *C. elegans*, deletion of endogenous nucleolar RG repeats leads to a loss of sub-nucleolar compartmentalization (Figure 4B) (Spaulding et al., 2022). How do RG repeats direct sub-nucleolar organization? Repeats may form interactions with proteins and/or RNA in one sub-compartment, thereby enriching the protein in that compartment and excluding it from others. If this were the case, we would expect full-length repeats to be capable of independently forming those interactions and enriching in the correct sub-nucleolar compartment. To the contrary, when ectopically expressed in *Xenopus* oocytes, RG repeats from Nucleolin and Fibrillarin do not enrich in a specific nucleolar sub-domain (Lavering et al., 2023). Our study confirms these results in *C. elegans*. One potential limitation of this study comes from the lower expression of NUCL-1 and FIB-1 RG repeats. Although it is challenging to precisely control expression levels in a living animal, studying RG repeats in their natural context is crucial for determining physiologically relevant functions. As demonstrated through visualization of endogenous GARR-1, concentration thresholds for sub-nucleolar phase separation have been met and nucleoli are correctly compartmentalized. Thus, regardless of over-expression levels, RG repeats should be free to enrich in pre-existing sub-compartments if they are capable. Instead, our study provides additional evidence from a living animal that full-length RG repeats do not independently enrich in sub-nucleolar compartments even when endogenous proteins are present and correctly partitioned (Figure 4C).

**Figure 4:**
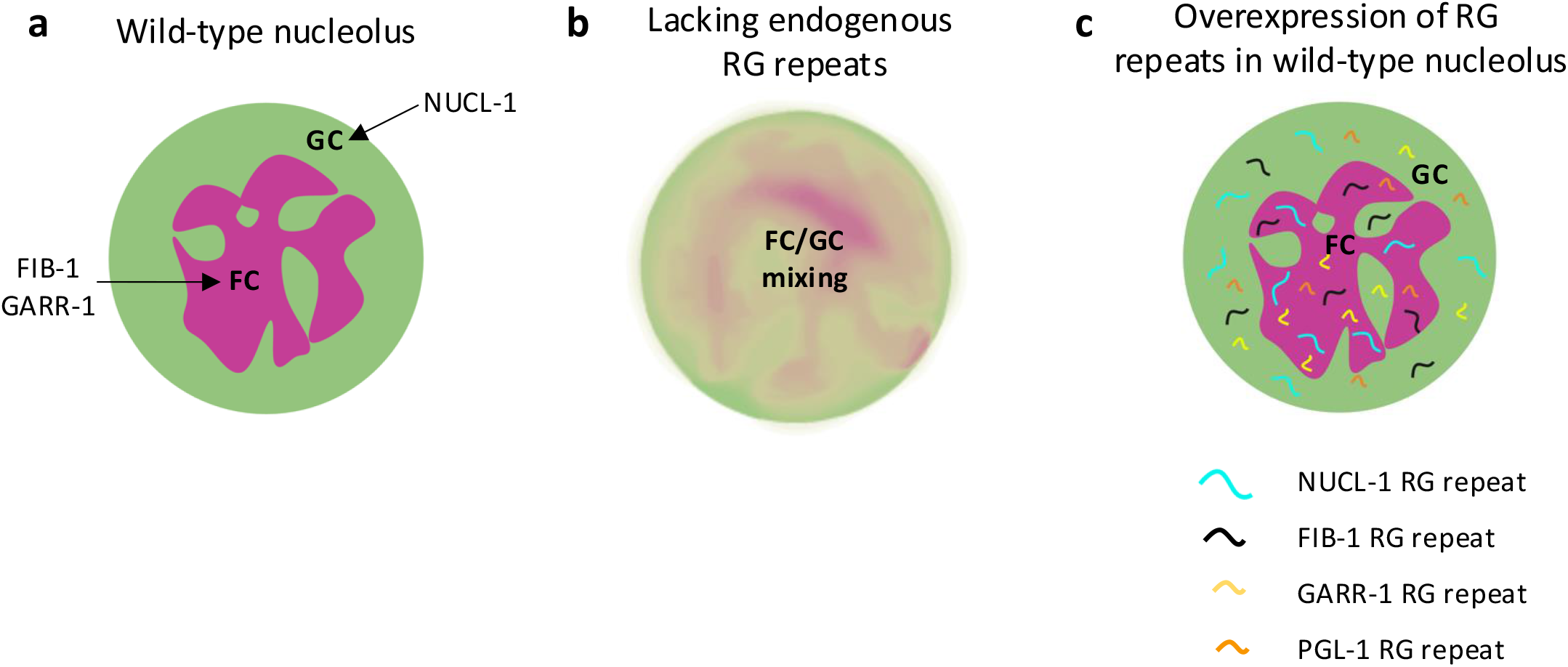
Model of RG repeat function in the *C. elegans* nucleolus. **a** In a WT nucleolus, binding of nucleolar proteins to functional partners may drive enrichment in a sub-compartment, while RG repeats allow for large-scale assembly of compartments. **b** When endogenous RG repeats are not present, nucleolar proteins do not compartmentalize but may still bind functional partners. **c** RG repeats accumulate within nucleoli but do not enrich in a sub-nucleolar compartment.

Our findings support a model in which RG repeats indirectly influence nucleolar organization by creating an environment conducive to compartmentalization that is primarily driven by other functional protein domains. For example, the methyltransferase domain of human FBL binds to nascent pre-rRNA and enriches FBL in the correct sub-nucleolar compartment. The FBL RG repeat controls larger-scale self-association (Yao et al., 2019). Recent work by King et al, also points to the role of functional RNA and DNA binding domains tethered to acidic domains (D/E tracts) as determinative factors in organization of nucleolar proteins (King et al., 2024).

*C. elegans* NUCL-1, FIB-1, and GARR-1 RG repeats are insufficient for sub-nucleolar compartmentalization, perhaps because they do not bind functional partners. When endogenous RG repeats from NUCL-1 or GARR-1 are deleted (NUCL-1ΔRG or GARR-1ΔRG), large-scale sub-nucleolar compartmentalization is lost. Surprisingly, NUCL-1ΔRG worms are healthy and fertile and GARR-1ΔRG worms show only mild fertility defects (Spaulding et al., 2022). This resilience may be restricted to organisms with a 2-compartment nucleolus and lost in animals with more complex nucleolar organization. In *C. elegans*, NUCL-1ΔRG and GARR-1ΔRG may still bind functional partners and take part in ribosome biogenesis despite the collapse of large-scale partitioning (Figure 4B). In summary, our data support the idea that sub-nucleolar organization is primarily driven by the functional interactions of structured protein domains, while RG repeats contribute to large-scale assembly of compartments by influencing the biophysical properties of nucleoli through additional specific or nonspecific interactions (Protter et al.) (Figure 4C).

Our previous work demonstrated that repeats are dispensable for nucleolar accumulation of endogenous proteins (Spaulding et al., 2022). If RG repeats are not required for nucleolar accumulation and are insufficient for sub-nucleolar compartmentalization, what is the significance of RG nucleolar and germ granule consensus motifs? In this study, nucleolar repeats with and without the phenylalanine-rich motif accumulated within nucleoli. The germ granule PGL-1 RG repeat contains a tyrosine-rich motif and also accumulates within nucleoli. Molecular interactions formed by RG repeats will vary in strength depending upon the identity and number of aromatic residues involved (Lin et al., 2017). Thus, distinctive consensus motifs may impart nucleolar and germ granule RG repeats with individual characteristics that help tune the biophysical conditions of each BMC to its function.

Defining the functions of IDRs inside a living animal is a crucial step in determining how mutations in these domains alter BMC dynamics and drive human disease (Tsang et al., 2020; Borcherds et al., 2021; Mensah et al., 2023). Our *in vivo* work supports a modulatory role of RG repeat domains in condensate organization and points to functional interactions as the primary drivers of nucleolar sub-compartmentalization (Choi et al., 2020; Savojardo et al., 2020; Feng et al., 2021). Extensive RG repeats like those found in NUCL-1, FIB-1, GARR-1, and their human homologs may act as flexible scaffolds, creating a nucleolar environment condusive to large scale compartmentalization. Disease-linked mutations in RG repeats likely disrupt the finely tuned biophysical properites of BMCs and lead to widespread functional consequences (Sheikh et al., 2022).

## METHODS

### Strain generation and maintenance

*C. elegans* strains were maintained using standard protocols (Brenner 1974). Strains created for this study include **DUP277** *glh-1(sam168[glh-1::T2A::sGFP2(1-10)::M3::NUCL-1RGG+NLS]) I*; **DUP281** *glh-1(sam171[glh-1::T2A::sGFP2(1-10)::M3::FIB-1RGG+NLS]) I*; **DUP282** *glh-1(sam172[glh-1::T2A::sGFP2(1-10)::M3::PGL-1RGG+NLS]) I*; **DUP284** *glh-1(sam174[glh-1::T2A::sGFP2(1-10)::M3::GARR-1NtermRGG+NLS]) I*; **DUP293** *garr-1(sam179[garr-1::wrmScarlet(1-11)]) IV*; **DUP294** *glh-1(sam168[glh-1::T2A::sGFP2(1-10)::M3::NUCL-1RGG+NLS]) I; garr-1(sam179[garr-1::wrmScarlet(1-11)]) IV*; **DUP295** *glh-1(sam171[glh-1::T2A::sGFP2(1-10)::M3::FIB-1RGG+NLS]) I; garr-1(sam179[garr-1::wrmScarlet(1-11)]) IV*; **DUP296** *glh-1(sam172[glh-1::T2A::sGFP2(1-10)::M3::PGL-1RGG+NLS]) I; garr-1(sam179[garr-1::wrmScarlet(1-11)]) IV*; **DUP297** *glh-1(sam174[glh-1::T2A::sGFP2(1-10)::M3::GARR-1NtermRGG+NLS]) I; garr-1(sam179[garr-1::wrmScarlet(1-11)]) IV*; **DUP305** *glh-1 (sam182[glh-1::T2A::sGFP2(1-11)::NLS]) I*; **DUP307** *glh-1 (sam182[glh-1::T2A::sGFP2(1-11)::NLS])1; garr-1(sam179[garr-1::wrmScarlet(1-11)]) IV*; **DUP311** *garr-1(sam179[garr-1::wrmScarlet(1-11)]) IV;nucl-1(sam186[nucl-1::sGFP(1-11)])IV*. Sequence files for CRISPR-generated alleles are stored on figshare (see Data Availability Statement). All strains generated for this study and their associated sequence files are available upon request.

### CRISPR strain construction

CRISPR/Cas9 genome editing was used to place a split superfolder-GFP11 tag (M3), a flexible linker sequence, a full-length RG sequence, and an SV40 nuclear localization signal on the C terminus of sGFP1(1-10) in the DUP223 background (GLH-1::T2A::sGFP2(1-10)). Creation of the DUP223 *glh-1(sam129[glh-1::T2A::sGFP2(1-10)]) I* allele was previously described (Goudeau *et al*., 2021). The 176 amino acid NUCL-1 RG repeat was inserted to create DUP277, the 107 amino acid FIB-1 RG repeat was inserted to create DUP281, the 48 amino acid long N-terminal GARR-1 RG repeat was inserted to create DUP284, and the 53 amino acid PGL-1 RG repeat was inserted to create DUP282. DUP305 was created by inserting a split superfolder-GFP11 tag, flexible linker, and SV40 nuclear localization signal on the C terminus of sGFP(1-10) in the DUP223 background. DUP293 was created by injecting GARR-1::wrmScarlet CRISPR constructs into the N2 laboratory strain. DUP294, DUP295, DUP296, DUP297, and DUP307 were created by crossing DUP277, DUP281, DUP282, DUP284, and DUP305 into DUP293, respectively. DUP311 was created by inserting sGFP onto the C-terminus of NUCL-1 in DUP293. CRISPR techniques for efficient genome editing in *C. elegans* were followed as described (Ghanta *et al*., 2021). All CRISPR reagents (Cas9 (Cat# 1081058), trRNA (Cat# 1072532), crRNAs (2nmol), and dsDNA repair templates (HDR donor blocks)) were ordered from Integrated DNA Technologies, Inc (San Diego, CA). Sequences for the guide RNA and repair templates are stored on figshare (https://figshare.com/articles/dataset/CRISPR_reagents_for_Spaulding_Updike_2024/26662102?file=48495868). All edits generated for this study were sequence verified, and sequence files are stored on figshare (https://figshare.com/articles/figure/Sequence_files_for_Spaulding_Updike_2024/26789935). All strains generated for this study are available upon request.

### Nucleolar imaging and analysis

L4 worms were plated at 20^°^C the day prior to imaging. On the day of imaging live, young adult worms were mounted on agarose pads in egg buffer (25mM HEPES (Fisher, cat#BP310-1), 120mM NaCl (Sigma, cat#S9888, 2mM MgCl_2_ (Sigma, cat#M9272, 50mM KCl (Fisher, cat#S77375-1, and 10mM levamisole (Thermofisher, cat# AC187870100) between the slide and a No.1.5 coverslip (Fisherbrand). Images were acquired using a point scanning confocal unit (LSM 980, Carl Zeiss Microscopy, Germany) on a Zeiss Axio Examiner Z1 upright microscope stand (ref: 409000-9752-000, Carl Zeiss Microscopy, Germany) equipped with a Plan-Apochromat 63X/1.4 Oil (ref:420782-9900-799, Carl Zeiss Microscopy, Germany) objective. sGFP and wrmScarlet fluorescence were excited with the 488nm Diode (0.5% laser power) and the 561nm DPSS laser (0.2% power), respectively. Fluorescence was collected with Airyscan2 with the following detection wavelengths: sGFP from 499 to 557nm and wrmScarlet from 573-627nm. Images of the adult germline were acquired using standard confocal mode. Images of pachytene nucleoli were sequentially acquired in Super Resolution mode (SR) at zoom 10, with a line average of 1, a resolution of 292×292 pixels, 0.043 x 0.043 μm pixel size, a pixel time of 0.69μs, in 16-bit, and in bidirectional mode. Z-stack images were collected with a step size of 0.170μm with the Motorized Scanning Stage 130×85 PIEZO (Carl Zeiss Microscopy) mounted on Z-piezo stage insert WSB500 (Carl Zeiss Microscopy). Microscope was controlled using Zen Blue Software (Zen Pro 3.1), Airy scan images were processed using the “auto” mode and saved in CZI format. Z stacks of at least 10 pachytene germ cell nucleoli were taken from each of 10 worms per genotype during at least 2 separate experiments.

ImageJ/Fiji was used to quantify the coefficient of variation (CV) within individual nucleoli. A single plane of the Z stack was chosen for each nucleolus that contained the maximum nucleolar area. A circle was placed within individual nucleoli that covered its maximum area without including background space and the following macro code was used to calculate the coefficient of variation (standard-deviation divided by the mean fluorescence):

getRawStatistics(N, mean, min, max, std);

print(std / mean);

The CV was measured from at least 10 nucleoli from each of 10 worms per genotype. Data was analyzed including all individual nucleoli or the mean of all nucleoli measured per worm.

ImageJ/Fiji was used to quantify the nucleolar:nucleoplasmic fluorescence intensity ratio of at least 50 nucleoli from at least 5 worms per strain using the same images as used for CV analysis. A single plane of the Z stack was chosen for each nucleolus that contained the maximum nucleolar area. A circle of the same size was used to measure the mean fluorescence intensity of 3 spots within the nucleolus, the nucleoplasm, and the surrounding cytoplasm. This was performed for both GFP and wrmScarlet. The average of the 3 cytoplasmic intensity spots was subtracted from the average of the 3 nucleolar and nucleoplasmic intensity spots. The background-subtracted nucleolar intensity was divided by the background-subtracted nucleoplasmic intensity to determine the ratio. Data was analyzed including all individual nucleoli or the mean of all nucleoli measured per worm.

Fiji was also used to create fluorescence intensity profile plots of pachytene germ cells using the same images as used for CV and nucleolar enrichment analysis. Dual-channel images were split into 2 images and the frames were synchronized. Brightness was auto-scaled for both GFP and wrmScarlet channels. A single plane of the Z stack was chosen for each nucleolus that contained the maximum cell area. A 6-micron straight line was placed across the center of a single cell in the GFP channel. The “plot profile” feature was used to create a plot of gray value vs distance in microns. The data was saved in the list format and imported into Prism. This was repeated for the wrmScarlet channel. Profile plots were created for at least 50 cells from at least 5 worms per strain.

Fiji was used to measure the Mander’s Colocalization Coefficient using the same images used for CV and profile plot measurements. Dual-channel images were split into 2 images and the frames were synchronized. A single plane of the Z stack was chosen for each image that contained the maximum number of nucleoli. The JACoP plugin was used to manually threshold the wSc and GFP channels and measure M1 and M2.

### Worm Crosses

Males were generated by plating 10 L4 hermaphrodites each on 10 plates and incubating at 30^°^C for 6 hours. Plates were then shifted to 20^°^C and males were picked 3 days later. DUP277, DUP281, DUP282, DUP284, and DUP305 males were crossed into DUP293 hermaphrodites. 4 F1 worms were picked from plates with 50% males (indicating successful mating). 9 F2 worms with both GFP and wrmScarlet expression were picked from each F1 clone using a fluorescent dissecting microscope. Homozygosity of F2 worms was determined by visually screening their progeny.

### Western Blotting

100-150ul of worms of each strain were washed off plates with dH_2_O and flash frozen in liquid nitrogen. 100ul of solubilization buffer (300mM NaCl, 50mM Tris-HCl [pH8.0], 10mM MgCl_2_, 1mM EGTA, ½ tablet Complete protease inhibitor, 1% Triton-X, 1mM PMSF) was added to each frozen worm pellet and worms were homogenized on ice for 1-2 minutes using a Fisherbrand cordless mixer with disposable pestle (Kimble, item#6CJ2ZNZ). Homogenate was left on ice for 1 hour and vortexed every 10 minutes, followed by centrifugation at 12K for 5 minutes at 4^°^C. The aqueous layer was transferred to a new tube and mixed with 1X Laemmli buffer (10% beta-mercaptoethanol (Fisher, cat#BP176-100), 4% SDS (Bio-Rad, cat#161-0203), 20% glycerol (Invitrogen, cat#15514-011), .004% bromophenol blue (Sigma, cat#B5525), 0.125M Tris-Cl pH6.8 (Fisher, cat#BP153-500)). Samples were then boiled for 10 minutes and spun at 12K for 5 minutes at room temperature. 30ug of protein as determined by Bradford protein quantification assay was loaded onto a mini-PROTEAN TGX stain-free gel (Bio-Rad, cat#4568084) and run at 200V for 25 minutes in SDS running buffer (.2501M Tris base (Fisher, cat#BP15201), 1.924M glycine (Fisher, cat#G48-500), .0347M SDS (Bio-Rad, cat#161-0203)). The gel was exposed to UV light for 5 minutes for total protein quantity detection and then the contents of the gel were transferred using a trans-blot turbo transfer pack (Bio-Rad, cat#1704156) on the Bio-Rad Trans-Blot Turbo Transfer System. The PVDF membrane was imaged to determine total protein levels and then blocked in 5% nonfat milk in TBST (20mM Tris-HCl [pH7.4], 150mM NaCl, 0.1% Tween) at room temperature for 1 hour. The membrane was incubated overnight at 4^°^C with rabbit polyclonal anti-GFP, 1:2000 (Invitrogen, cat#A-6455) in 5% milk and then washed 6X 10 minutes in TBST at room temperature. The membrane was then incubated in goat anti-rabbit IgG-HRP, 1:10,000 (Bio-Rad, cat#170-6515) in 5% milk for 1 hour at room temperature and washed 6X 10 minutes in TBST. Finally, the membrane was developed using Clarity Western ECL Substrate (Bio-Rad, cat#1705060S) and imaged on a Syngene G:Box gel and blot imaging system. For images of uncropped blot see Source Data.

### Statistics and Reproducibility

All imaging experiments investigating nucleolar accumulation and sub-nucleolar organization (Figures 2,3; Supplemental Figures 2,3) were performed on at least two independent occasions and similar results were always obtained. Imaging and image analysis was not done blinded to genotype because it was performed sequentially as each strain was created. When imaging DUP311 worms we would always observe some worms with less precise FC/GC organization (approximately 25% of total imaged worms, observed at each imaging session). This phenomenon may indicate that the sGFP and wrmScarlet tags on NUCL-1 and GARR-1, respectively, are interfering with the stability of nucleolar substructure.

Worms with less precise FC/GC organization were still included in all analyses. Western blots were performed on three independent occasions and similar results were always obtained (Supplemental Figure 1). For all pairwise comparisons (Figures 2a-f,3a-f) unpaired, 2-tailed t tests with Welch’s correction was performed. For comparisons of three or more groups (Figures 2g-i,3g-i and Supplemental Figure 1c and 3) one-way ANOVA tests with multiple comparisons were performed. Statistical analysis was done using Prism software.

## DATA AVAILABILITY

For CRISPR/Cas9 editing experiments, sequences for the guide RNA and repair templates are stored on figshare (https://figshare.com/articles/dataset/CRISPR_reagents_for_Spaulding_Updike_2024/26662102?file=48495868). All edits generated for this study were sequence verified and sequence files are stored on figshare (https://figshare.com/articles/figure/Sequence_files_for_Spaulding_Updike_2024/26789935). All strains generated for this study are available upon request. Source data for all experiments are provided with the paper.

## ACKNOWLEDGEMENTS

We would like to acknowledge support from Chris Smith in the MDI Biological Laboratory (MDIBL) Sequencing Facility, Dr. Frederic Bonnet in the MDIBL Light Microscopy Facility, and plate pouring services provided by the MDIBL Animal Resources Core. Schematics created with BioRender.com.

## AUTHOR CONTRIBUTIONS

E.L.S. generated worm strains, performed imaging, data analysis, and experimental design, and wrote the manuscript. D.L.U. provided conceptualization and experimental design and edited the manuscript.

