## Supplemental Figures 1-4 for "*C. elegans* nucleolar RG repeats are sufficient for nucleolar accumulation but insufficient for sub-nucleolar compartmentalization"

### Supplemental Figure 1: RG repeats and their expression

**a**

| Protein name | Native BMC localization | Length of RG repeat | RG repeat amino acid sequence |
|---|---|---|---|
| NUCL-1 | nucleolus-GC | 176 amino acids | RGRGGGGFRGGRGGGSGFTPRGGGGGFRGGDRGRSPGGFRGGDRGRSSSGFRGGDRGRSPGGFRGDREGGFSPRGRGGGFRGGDRGGFRGGDRGGSPWRGGDRGNFRGGDRGGSPWRGGDRGVANRGRGDFSGGSRGGNKFSPRGGGRGGFTPRGRGGNDFSPRGGRGNFQSRGGG |
| FIB-1 | nucleolus-FC | 107 amino acids | RGGGGGGFRGGRGGDRGGSRGGFGGGGRGGYGGGDRGSFGGGDRGGFRGGRGGGDRGGFRGGRGGGDRGGFGGRGSPRGGFGGRGSPRGGRGSPRGGRGGAGGMRGG |
| GARR-1 | nucleolus-FC | 48 amino acids | RGGRGGGGGFRGGRGGGGGGGFRGGRGGDRGGGFRGGRGGFGGGGRGG |
| PGL-1 | germ granule | 53 amino acids | RGGYGGGDRGGRGGYGGDRGGRGGYGGGDRGGRGGYGGDRGRGGYGGRGGRGG |

**b**

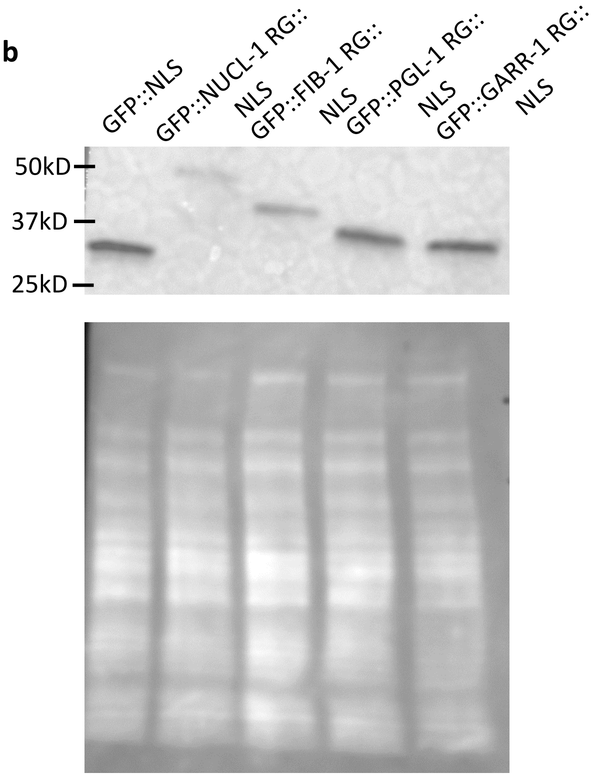


**c**

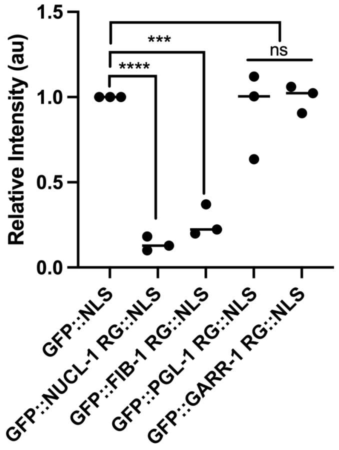


**a** Table including the RG repeats used in this study. Nucleolar and germ granule consensus motifs are underlined. Phenylalanine residues and motifs of nucleolar repeats are highlighted in purple while tyrosine residues and motifs from the germ granule repeat are highlighted in blue. **b** A representative Western blot against GFP confirms the expected sizes of CRISPR/Cas9-inserted RG repeats (top). Total protein (below) on the same blot. **c** Quantification of western blot GFP band intensities relative to total protein. Points on graph represent individual western blots with median shown. One-way ANOVA with multiple comparisons. ***p<.01, ****p>.001.

### Supplemental Figure 2: RG repeats accumulate in germline nucleoli

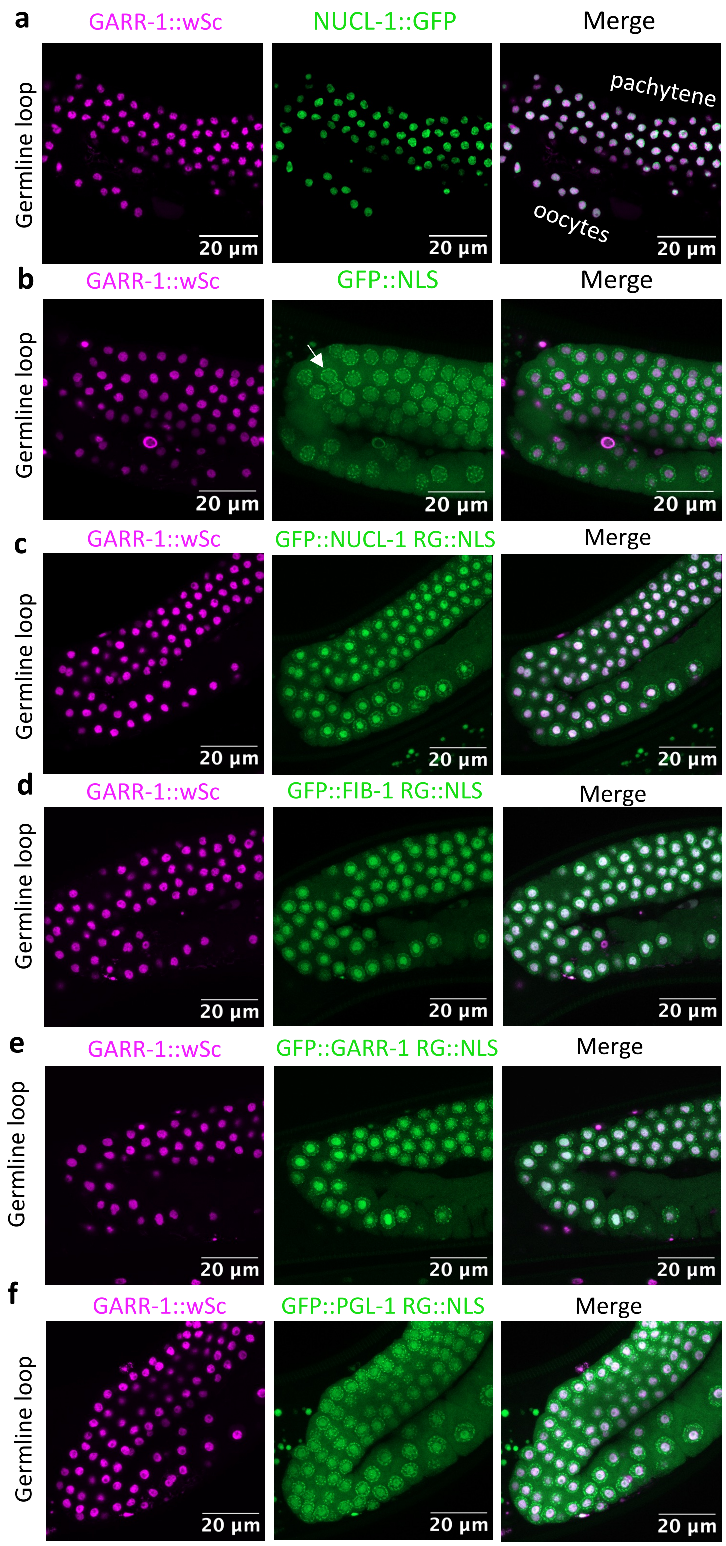


**a-f** Confocal images of adult hermaphrodite germline loop. **a** Endogenous GARR-1 and NUCL-1 are highly enriched in germ cell nucleoli. **b** GFP::NLS accumulates throughout nuclei but is only slightly enriched in nucleoli. **c-f** RG repeats from NUCL-1 (**c**), FIB-1 (**d**), GARR-1 (**e**), and PGL-1 (**f**) enrich in GARR-1-labeled nucleoli. All images are maximum projections of an entire z stack, except for **c**, which is a projection of 22 out of 30 slices to account for worm movement.

### Supplemental Figure 3: Mander's Colocalization Coefficient

**a**

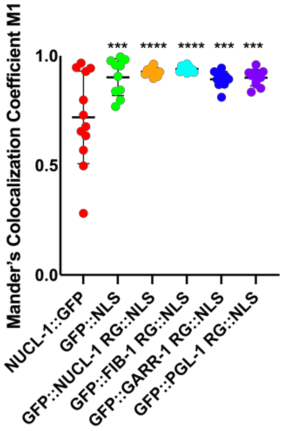


**a**The Fiji plugin, JACoP, was used to measure Mander's Overlap Coefficient 1 (M1). M1 reports on the percentage of GARR-1::wSc fluorescence in areas of GFP fluorescence. Each dot represents the mean M1 per worm. At least 2 images per worm were analyzed, with 2-5 nucleoli in each image. NUCL-1::GFP, n=12 worms; GFP::NLS, n=10 worms; GFP::NUCL-1RG::NLS, n=10 worms, GFP::FIB-1RG::NLS, n=10 worms, GFP::GARR-1RG::NLS, n=11 worms, GFP::PGL-1RG::NLS, n=11 worms. Lines represent the mean of M1 ± SD. One-way ANOVA with multiple comparisons. ***p<.001, ****p<.0001.

### Supplemental Figure 4: Mean fluorescence intensity traces

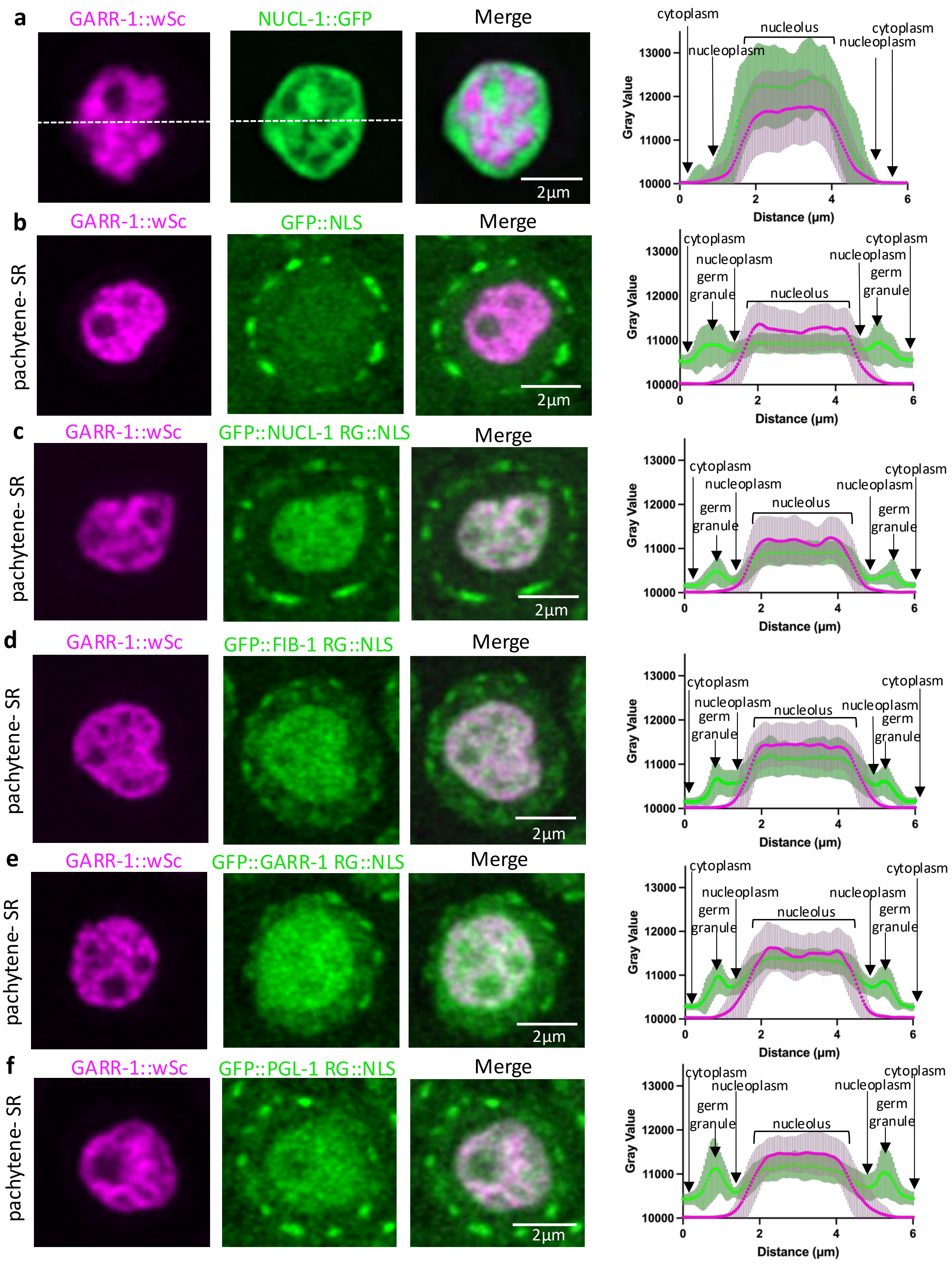


**a-f** A 6-micron line (white dashed line) was drawn through the center of at least 50 pachytene germ cells per strain to measure the fluorescence intensity of the cytoplasm, nuclear envelope (including germ granules), nucleoplasm, and nucleoli. Within single cells, the line was positioned in the same location for GFP and wrmScarlet channels. Traces were used to create plots of mean fluorescence intensity. **a** n=55 germ cells from 6 worms. **b-f** n=50 germ cells from 5 worms. Lines represent the mean fluorescence ± SD from 50-55 germ cells.
